# Enhanced Working Memory Representations for Rare Events

**DOI:** 10.1101/2024.03.20.585952

**Authors:** Carlos Daniel Carrasco, Aaron Matthew Simmons, John E. Kiat, Steven J. Luck

## Abstract

Rare events (*oddballs*) produce a variety of enhanced physiological responses relative to frequent events (*standards*), including the P3b component of the event-related potential (ERP) waveform. Previous research has suggested that the P3b is related to working memory, which implies that working memory representations will be enhanced for rare stimuli. To test this hypothesis, we devised a modified oddball paradigm where a target disk was presented at one of 16 different locations, which were divided into rare and frequent sets. Participants made a binary response on each trial to report whether the target appeared in the rare set or the frequent set. As expected, the P3b was much larger for stimuli appearing at a location within the rare set. We also included occasional probe trials in which the subject reported the exact location of the target. Accuracy was higher for rare than frequent locations. In addition, memory reports on rare trials were more accurate in participants with larger P3b amplitudes on rare trials (although reports were not more accurate for trials with larger P3b amplitudes within participants). We also applied multivariate pattern analysis to the ERP data to “decode” the remembered location of the target. Decoding accuracy was greater for locations within the rare set than for locations within the frequent set. We then replicated and extended our behavioral findings in a follow-up experiment. These behavioral and electrophysiological results demonstrate that although both frequent and rare events are stored in working memory, working memory performance is enhanced for rare oddball events.

**Impact Statement:** For many decades, researchers have observed that rare events elicit a broad range of physiological responses, and there has been much speculation about the functional significance of these responses. One such response is the P3b component, which is a large voltage deflection in scalp EEG recordings. Over 40 years ago, the P3b was hypothesized to reflect “context updating” (now often called “working memory updating”). However, there has been no direct evidence that working memory is actually enhanced for rare, P3b-eliciting events. In the present study, we found that both behavioral and electrophysiological measures of working memory were enhanced for rare events. However, it is not clear that the increased P3b-related brain activity actually caused the enhancement of working memory.

## Introduction

Rare task-relevant events (often called *oddballs*) generate a variety of physiological responses, including increased firing of noradrenergic neurons in the locus coeruleus, phasic pupil dilation, increased blood-oxygen-level dependent (BOLD) activity in a variety of cortical areas, and a large P3b event-related potential (ERP) component (Bledowski et al., 2004; Clark et al., 2000; Johnson, 1988; Krebs et al., 2018; Linden et al., 1999; Murphy et al., 2011; Polich, 1986; Soltani & Knight, 2000). There has been considerable speculation about the functional significance of these physiological changes (Kim, 2014; Linden et al., 1999; Nieuwenhuis et al., 2011; Paller et al., 1992). A common hypothesis is that the P3b activity elicited by oddballs is related to working memory encoding, although the details vary across theories (Donchin & Coles, 1988; Kok, 2001; Polich, 2007, 2012). Moreover, oddballs elicit phasic increases in attention (Aston-Jones & Cohen, 2005; Katayama & Polich, 1998; Kim, 2014; Murphy et al., 2011), which might also enhance working memory. However, we know of no direct evidence that working memory is actually enhanced for relatively rare events compared to relatively frequent events. The goal of the present study was to test this hypothesis, using both behavioral and electrophysiological measures.

Typical oddball paradigms do not provide a sensitive assessment of working memory. For example, a typical paradigm would involve presenting a sequence of stimuli in which 90% are the letter X and 10% are the letter O, and the task would be to press one button for Xs and another button for Os. The responses are made immediately, so it is not necessary to store the Xs and Os in working memory. Moreover, the Xs and Os are so easily discriminable that memory performance would likely be at ceiling if tested after a brief delay.

We therefore developed the modified oddball paradigm shown in Figure 1. On each trial, a target disc appeared briefly near one of eight locations around a circle, and the main task was to press one button if the disc appeared near one of the cardinal axes (up, down, left, or right) and a different button if it appeared near one of the diagonal axes (upper left, upper right, lower left, or lower right). One of these two categories was rare (12.5%), and the other was frequent (87.5%). This was much like a traditional oddball task, in which participants make an immediate response to indicate whether the stimulus belonged to the rare category or the frequent category. We assumed that stimuli belonging to the rare^1^ category (the oddballs) would elicit a larger P3b component than stimuli belonging to the frequent category (the *standards*). To provide a sensitive measure of the working memory representation of the disk, we also included occasional *probe* trials, on which participants were asked to click on the exact location of the disc from that trial after a brief delay. If working memory is enhanced for rare stimuli, then the target localization response on probe trials should be more accurate following an oddball than following a standard. Moreover, if the P3b component elicited by oddballs is associated with working memory updating, then participants with greater P3b amplitudes for the oddballs should exhibit more accurate memory on the probe trials than individuals with smaller P3b amplitudes.

**Figure 1.**
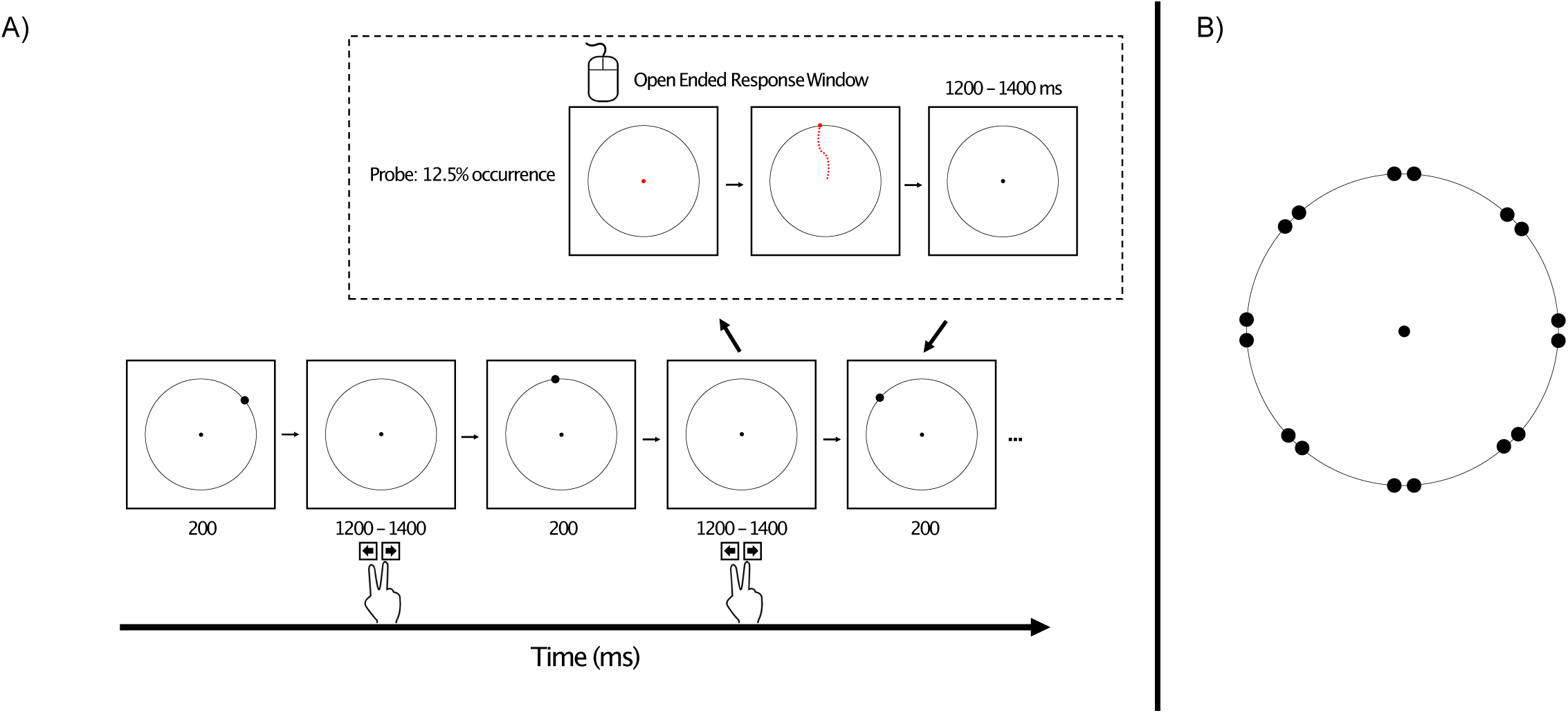
A) Overview of the oddball task. On every trial, participants made a speeded keypress following the presentation of a target disc to indicate whether it appeared near a cardinal axis or near a diagonal axis (as shown in B). For a given trial block, one axis was rare, and the other was frequent. On 12.5% of trials (probe trials), the fixation point turned red at the offset of the intertrial interval, which signaled participants that they should use the mouse to click on the remembered location of the target disk from that trial. (B) Possible locations of the target disc, which could appear 1° counterclockwise or 1° clockwise from a cardinal or diagonal axis. Note that the distance of the targets from the axes is slightly exaggerated in this image.

In addition, we applied multivariate pattern analysis (MVPA) to the ERP data to *decode* the remembered location of the disc in the delay period following each stimulus. This provided a means of monitoring the working memory representation of the stimuli during the delay period following each target, independently of the decision and response processes that are involved in the behavioral responses. Specifically, we decoded which of the four locations within a category was presented (e.g., up, down, left, or right for the cardinal category). We predicted that the within-category decoding accuracy would be greater for a given category when that category was rare than when it was frequent.

## Experiment 1

### Methods

#### Participants

Twenty-two human participants from the UC Davis community completed the study (14 women, 7 men, 1 unreported gender; mean age = 20, SD = 1.69). All participants were neurotypical and had normal or corrected-to-normal vision with no history of neurological conditions. Twenty participants were right-handed, and two were left-handed. We chose to collect a somewhat higher N than is typical of multivariate pattern analysis EEG studies (e.g., N = 16 in Bae & Luck, 2019; Bae, 2021; Bae & Luck, 2018) because of the relatively small number of oddball trials in the present study. Monetary compensation was provided at a rate of $15/hr. All participants provided informed consent, and the study was approved by the University of California, Davis Institutional Review Board.

#### Stimuli and Task

Figure 1 illustrates the stimuli and task. The experiment presentation script, data, and analysis scripts can be downloaded at <u>doi.org/10.17605/OSF.IO/HV7JU</u>.

Stimuli were presented using Psychopy (Peirce et al., 2019) on an LCD monitor (Dell U2412M) with a gray background (27.7 cd/m^2^) at a viewing distance of 100 cm. A 0.05° black fixation dot was continuously visible in the center of the monitor, surrounded by a black circle with a radius of 2.17°.

On each trial, a black target disc (0.2°) appeared for 200 ms, followed by an interstimulus interval of 1200-1400 ms (rectangular distribution). The disk appeared on the circle, near one of the cardinal axes (0°, 90°, 180°, 270°) or near one of the diagonal axes (45°, 135°, 225°, 315°), centered either ±1° from one of these axes (e.g., 44°, 46°, 89°, 91°). Thus, there were 16 possible target locations, 8 near the cardinal axes and 8 near the diagonal axes. Placing the target ±1° from an axis was designed to require participants to remember the precise location of the target rather than relying on simple categories such as “top” or “lower left”. In addition, this made it possible to examine biases away from the cardinal axes (Bae, 2022).

For four of the eight trial blocks, the target appeared near a cardinal axis on 12.5% of trials (oddballs) and appeared near a diagonal axis on the remaining 87.5% of trials (standards). This was reversed for the other four trial blocks. Each of the eight locations within the cardinal or diagonal category occurred with equal probability. Participants were instructed to make a speeded response on a computer keyboard to indicate whether the immediately preceding target was near a cardinal axis or near a diagonal axis. Half the participants pressed the left arrow key for cardinal and the right arrow key for diagonal; this was reversed for the other half.

Although most oddball studies use only one rare stimulus (e.g., a high-pitched tone) and one frequent stimulus (e.g., a low-pitched tone), the P3b component is sensitive to the probability of the task-defined category rather than the probability of the physical stimulus (Luck, 2014; Mecklinger & Ullsperger, 1993). Oddball studies have used abstract categories such as the ones used here for decades (e.g., Kutas et al., 1977).

Each block contained 256 trials (32 oddballs and 224 standards). The blocks alternated between cardinal-oddball/diagonal-standard and diagonal-oddball/cardinal-standard, with the starting condition counterbalanced across participants. Each participant received 256 oddballs and 1792 standards, with each of the 16 locations occurring equally often within each of these categories (2048 trials total). Thus, there were 112 trials for a given location when that location was in the standard category and 16 trials for a given location when that location was in the oddball category.

Working memory for the exact target location was probed following a random 12.5% of targets. When a target was probed, the fixation dot changed from black to red at the end of the interstimulus interval (i.e., after the participant indicated whether the target had been near a cardinal or diagonal axis). Once it turned red, the participant could use the mouse to move the red dot. They were instructed to move the red dot to the remembered location of the target and then click the mouse button (with no time pressure). The red dot then disappeared and was replaced by the black fixation dot in the center of the monitor. The stream of targets then resumed after a delay of 1200-1400 ms. For each participant, oddballs were probed on 32 trials (4 probes per block) and standards were probed on 224 trials (28 probes per block).

Note that participants could not predict whether a given trial would be a probe trial. Probe trials were just like any other trials until the end of the intertrial interval following a given target. Thus, the working memory representations and neural activity during the intertrial interval following a target could not be systematically different on probe and non-probe trials.

We probed on only a small subset of trials for two reasons. First, we wanted the task to be more like a traditional oddball task, in which exact memories are not probed. Second, probe responses took considerable time, and we could obtain more targets in a session of a reasonable duration if we probed infrequently. A very large overall number of trials was needed to obtain a sufficient number of trials per location for the EEG decoding (which used the data from all trials, not just probe trials). Far fewer trials were needed to obtain robust measures of behavioral accuracy.

#### EEG Recording and Preprocessing

The continuous EEG was recorded using a Brain Products actiCHamp recording system. We recorded EEG signals from 27 standard 10/20 sites: FP1, Fz, F3, F7, Cz, C3, Pz, P3, P5, P7, P9, PO7, PO3, O1, POz, Oz, FP2, F4, F8, C4, P4, P6, P8, P10, PO4, PO8, O2. We also recorded signals from horizontal electrooculogram (HEOG) electrodes lateral to the left and right external canthi, from a vertical electrooculogram (VEOG) electrode located under the right eye, and from electrodes over the left and right mastoids. Single-ended voltages were recorded relative to a ground electrode located at AFz. All electrode impedances were kept < 50 KΩ. The signals were filtered with a cascaded integrator-comb antialiasing filter (half-power cutoff at 130 Hz) and digitized at 500 Hz.

All preprocessing steps were conducted in MATLAB using the EEGLAB and ERPLAB toolboxes (Delorme & Makeig, 2004; Lopez-Calderon & Luck, 2014), following the standard pipeline described by Luck (2023).We began by shifting the event codes to account for the monitor delay (56 ms, which was measured using a photodiode). The signals were then resampled at 250 Hz. The DC offset was removed, and the signals were high-pass filtered (noncausal Butterworth impulse response function, half-amplitude cutoffs at 0.1 Hz, 12 dB/oct roll-off). Time segments between the trial blocks were deleted, and the EEG data were referenced to the P9 electrode site. A bipolar VEOG channel was created by subtracting the FP2 electrode from the VEOG electrode, and a bipolar HEOG channel was created by subtracting HEOG-right from HEOG-left.

Independent component analysis (ICA) was then performed to correct for blinks and eye movements (excluding the bipolar channels). The data used for the ICA decomposition were filtered more aggressively (noncausal Butterworth impulse response function, half-amplitude cutoffs at 1 – 30 Hz, 48 dB/oct roll-off) and resampled at 100 Hz. The ICA weights were then transferred back to the original data, and independent components corresponding to blinks and artifacts were removed from the data; typically, 1-2 components were removed per participant. We used consistency between the shape, timing and spatial location of the component compared to the bipolar HEOG and VEOG signals to determine which components were artifacts.

The ICA-corrected data were then re-referenced to the average of the left and right mastoid electrodes. The data were then segmented from -500 to 1496 ms relative to stimulus onset and baseline-corrected to the mean voltage from -500 to 0 ms. Finally, epochs were marked for rejection using standard ERPLAB routines if they contained large voltage deflections in any channel (using a simple voltage threshold) or an eyeblink that would have prevented perception of the target (using a moving window peak-to-peak function from 0 to 200 ms in the bipolar VEOG channel). This led to an average rejection of 13.7% of trials across participants (SE = 6.4%). We always exclude participants for whom more than 25% of trials were rejected; no participants exceeded this threshold in the present study.

#### Behavioral Analyses

For the oddball categorization task, we computed the proportion correct and the mean response time (RT) for each participant, separately for the oddball and standard categories.

For the probe task, the stimuli and responses were coded in terms of their angular position around the circle of possible target locations. We excluded trials on which the reported location was > 40° away from the true location (0.89% ± 0.31% of trials), because such large errors presumably reflect lapses of attention. On the remaining trials, we computed the *response error*, defined as the angular distance between the true target location and the reported location. In the primary analyses, we took the absolute value of the response error on each trial and averaged across trials for a given participant, separately for the oddball and standard categories. As a secondary analysis, we took the standard deviation of the response errors (see supplementary Figure S1).

Working memory representations tend to be biased away from the cardinal axes (Bae, 2022) and we examined these biases in a separate analysis of the probe data. Following prior research, we expected that locations that were 1° clockwise from a cardinal axis would be reported as being more than 1° clockwise, and that locations that were 1° counterclockwise from a cardinal axis would be reported as being more than 1° counterclockwise. If working memory representations are enhanced for oddballs compared to standards, then these biases should be reduced for the oddballs. To analyze the biases, we took the response error on each trial and gave it a negative sign if it was farther away from the nearest cardinal axis than the true stimulus location (e.g., given a true location of 89°, which was near the cardinal axis at 90°, a reported location at 87° was coded as a response error of -2°). Similarly, we gave a response error a positive sign if it was biased in the opposite direction (e.g., given a true location of 89°, a reported location at 91° was coded as a response error of +2°). These values were then averaged across trials for a given participant, separately for oddball and standard trials. A consistent repulsion away from the nearest axis would lead to a negative average value. Little or no bias of this sort would be expected for locations near the diagonal axes for either oddballs or standards, so trials with stimuli near the diagonal axes were excluded from this analysis.

#### P3b Scoring

We measured P3b amplitude at an a priori electrode size (Pz) and an a priori time window which we defined as defined as ±150 ms around the P3b peak of the grand averaged ERP. This peak was at 512 ms so our measurement window was from 362 to 662 ms. The P3b amplitude was scored as the mean voltage during this time window at Pz, separately for oddballs and standards.

#### Decoding Analysis

The decoding analysis collapsed across the two locations that were ±1° from a given axis (e.g., 44° and 46°), which were too similar to be reliably differentiated by the decoding process. This gave us four cardinal locations (0°, 90°, 180°, 270°) and four diagonal locations (45°, 135°, 225°, 315°). We performed the decoding separately for these two categories, separately, when a category was the oddball and when it was the standard. For example, we decoded which of the four cardinal locations was present on the cardinal trials when the cardinal category was the oddball, and we separately decoded which of the four cardinal locations was present on the cardinal trials when the cardinal category was the standard. This allowed us to compare the decoding accuracy for the same stimulus locations when those locations were oddballs and when they were standards. Decoding was performed separately at each time point (every 4 ms from - 500 to +1496 ms).

We followed the location decoding procedure developed by Bae and Luck (2018), as implemented in ERPLAB Toolbox (Lopez-Calderon & Luck, 2014) using MATLAB’s fitcecoc() function. The first step was to apply a 6 Hz lowpass filter (48 dB/octave roll-off), which minimized contamination from alpha-band EEG oscillations. This filter reduced the temporal resolution of the analysis, but that was not a major problem given that we were examining long-duration working memory effects. The ocular channels were left out of the analysis, leaving 27 scalp channels.

Decoding was performed on averaged ERP waveforms using support vector machines (SVM) with error-correcting output codes (Dietterich & Bakiri, 1995) and 3-fold cross-validation. We conducted four separate decoding runs for each participant: a) cardinal oddball; b) cardinal standard; c) diagonal oddball; and d) diagonal standard. For each run, there were four possible stimulus locations (the four different cardinal locations or the four different diagonal locations). The decoder attempted to determine which of these four locations was presented on the basis of the pattern of voltage across electrode sites at a given time point.

Because decoding accuracy tends to increase when more trials are available, we randomly subsampled a subset of trials from the standards to equate the number of trials for oddballs and standards. This yielded an average across participants of 19.5 trials (SD = 4.5) at each of the four locations for each decoding run. The available trials were randomly divided into three subsets, and a separate averaged ERP waveform was created from each of these three subsets. This gave us a matrix of 4 classes (4 locations) ξ 3 averages ξ 27 electrode sites ξ 500 time points for each participant for each decoding run. Decoding was performed on the matrix of classes ξ averages ξ electrode sites, separately at each time point for each participant for a given run.

We performed a 3-fold cross-validation by training the decoder on two of the three averages for each location and then having the decoder predict the location on the basis of the remaining average for each location. We then repeated the process two more times with new decoders, changing which averages were used for training and which were used for testing. This process was then iterated 100 times, with different random subsets of trials used to create the averaged ERPs for each iteration. Completely new decoders were trained for every fold, iteration, time point, and participant.

For each fold, the decoder consisted of four separate SVMs, each of which was trained to distinguish between one location and the other three locations on the basis of the pattern of voltage across electrode sites. To test the decoder, the vector of voltages across electrode sites for a given location from the test average was passed to all four SVMs. The decoder predicted the location for that test average using MATLAB’s predict() function, which minimized the average binary loss over the four SVMs. Decoding accuracy was computed as the proportion of correct predictions by the decoder across the three folds and 100 iterations. Because there were four locations, chance decoding accuracy was 25%.

We then averaged decoding accuracy across the cardinal and diagonal decoding runs, separately for oddballs and standards. This gave us a decoding accuracy for each participant at each time point for oddballs and for standards. To maximize statistical power, decoding accuracy was then averaged across an a priori time window that began at the start of the P3b measurement window (362 ms) and extended until the end of the epoch (1496 ms). For each participant, this gave us one decoding accuracy value for oddballs and another for standards.

### Results

#### Behavioral results

##### Oddball task

We first examined the speed and accuracy for the buttonpress task, in which participants indicated whether a given target was an oddball or a standard. As is typical in oddball tasks, mean accuracy was lower for oddballs (80.5% ± 2.32%) than for standards (95.7% ± 0.512%), which was significant in a paired *t* test (*t*(21) = 6.85, *p* < 0.0001, *d_z_* = 1.46). In addition, mean RTs were slower for oddballs (470.66 ± 19.48 ms) than for standards (346.04 ± 18.75 ms), which was also a significant difference (*t*(21) = 14.59, *p* < 0.0001, *d_z_* = 3.11). Note that all statistical tests reported in this paper were two-tailed and used an alpha of .05. Effect sizes are quantified as d*_z_*, which is the standard effect size metric corresponding to paired and one-sample *t* tests (Cohen J, 1988; Lakens, 2013). When means are given, we also provide the standard error of the mean (SEM).

##### Probe task

We next examined accuracy on the probe trials, for which participants clicked on the remembered location of the target. Figure 2a shows that the mean absolute response error for the probe task was smaller for oddballs (2.99 ± 0.155°) than for standards (3.36 ± 0.156°), which was a significant difference in a paired *t* test (*t*(21) = 3.30, *p* = 0.003, *d_z_* = 0.704). The same pattern was found when the standard deviation of response errors was used as the metric of memory precision (supplementary Figure S1). These results indicate that working memory representations were more accurate for rare stimuli than for frequent stimuli.

**Figure 2.**
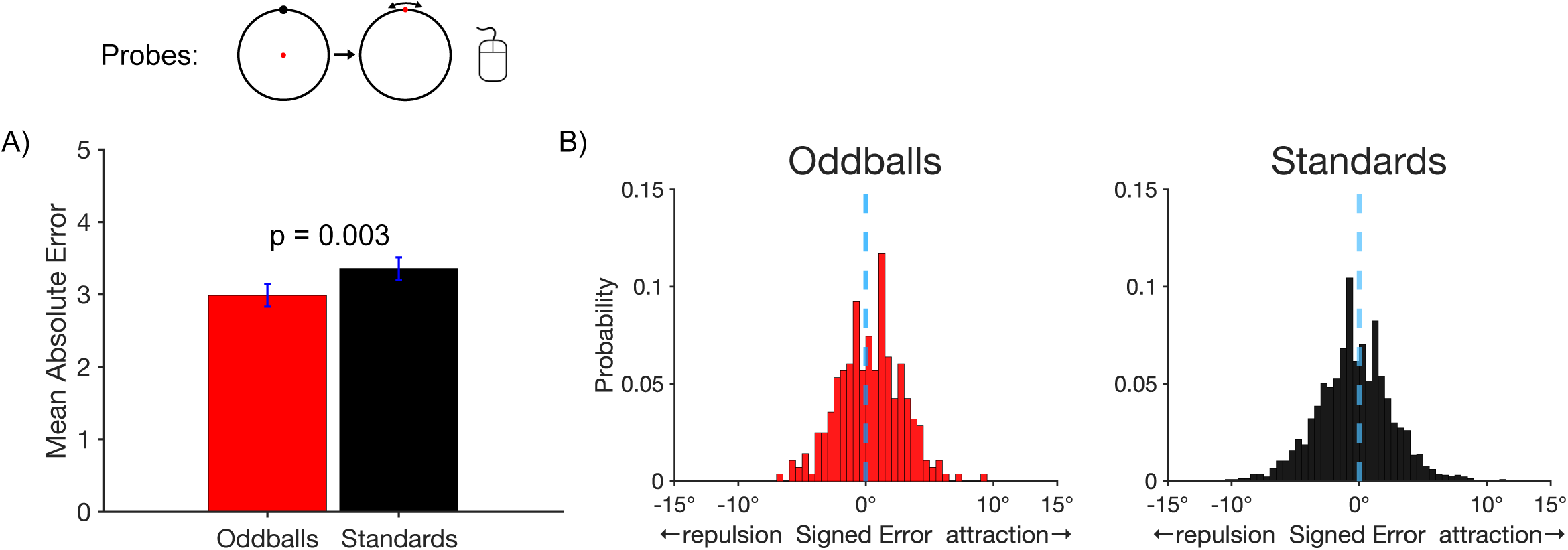
Probe trial results. A) Mean absolute error in the report of the exact location of the target disc. Error bars show ±1 SEM. B) Histograms of the single-trial bias values for target discs near a cardinal axis. For oddballs, the distribution of errors was fairly symmetrical around zero (minimal bias). For standards, the distribution is shifted toward negative values (repulsion away from the axis).

We also asked whether working memory representations of rare stimuli are less subject to systematic biases. In particular, we took advantage of the fact that stimuli presented near a cardinal axis are typically remembered as being shifted away from that axis (Bae, 2022; Pratte et al., 2017; Wei & Stocker, 2015). We asked whether this bias would be reduced for oddballs relative to standards. Figure 2b shows histograms of the single-trial bias values, with negative values indicating a bias away from the nearby cardinal axis and positive values indicating a bias toward the axis. On standard trials, the distribution of values was shifted toward the negative side of zero, indicating the typical finding of repulsion away from the axis. We computed an average bias score for each participant, and we found that this bias score was significantly different from zero for the standards in a one-sample *t* test (*t*(21) = 14.59, *p* = 0.002, *d_z_* = 0.739). On oddball trials, the distribution of bias values was more symmetrical around zero, and the average bias score was not significantly different from zero (*t*(21) = 1.08, *p* = 0.292, *d_z_* = 0.231). The key finding was that the average bias score was significantly more negative for standards than for oddballs in a paired *t* test (*t*(21) = 3.17, *p* = 0.005, *d_z_* = 0.676). This indicates that working memory representations were less biased for rare stimuli in addition to being more accurate.

#### P3b amplitude

Figure 3a shows the grand average ERP waveforms for oddballs and standards at five representative electrode sites. As is typical, a large P3b was visible for oddballs, peaking at 512 ms at the Pz electrode site. As shown in Figure 3b, the mean voltage during the measurement window (362-662 ms) at Pz was significantly larger for oddballs than for standards in a paired *t* test (*t*(21) = 8.10, *p* < 0.0001, *d_z_* = 1.43). The scalp distribution of the oddball-minus-standard difference wave during this window showed the typical midline centroparietal maximum (Figure 3c). Thus, although the oddball paradigm used in the present study was somewhat unusual, it yielded the typical pattern of a larger P3b for oddballs than for standards.

**Figure 3.**
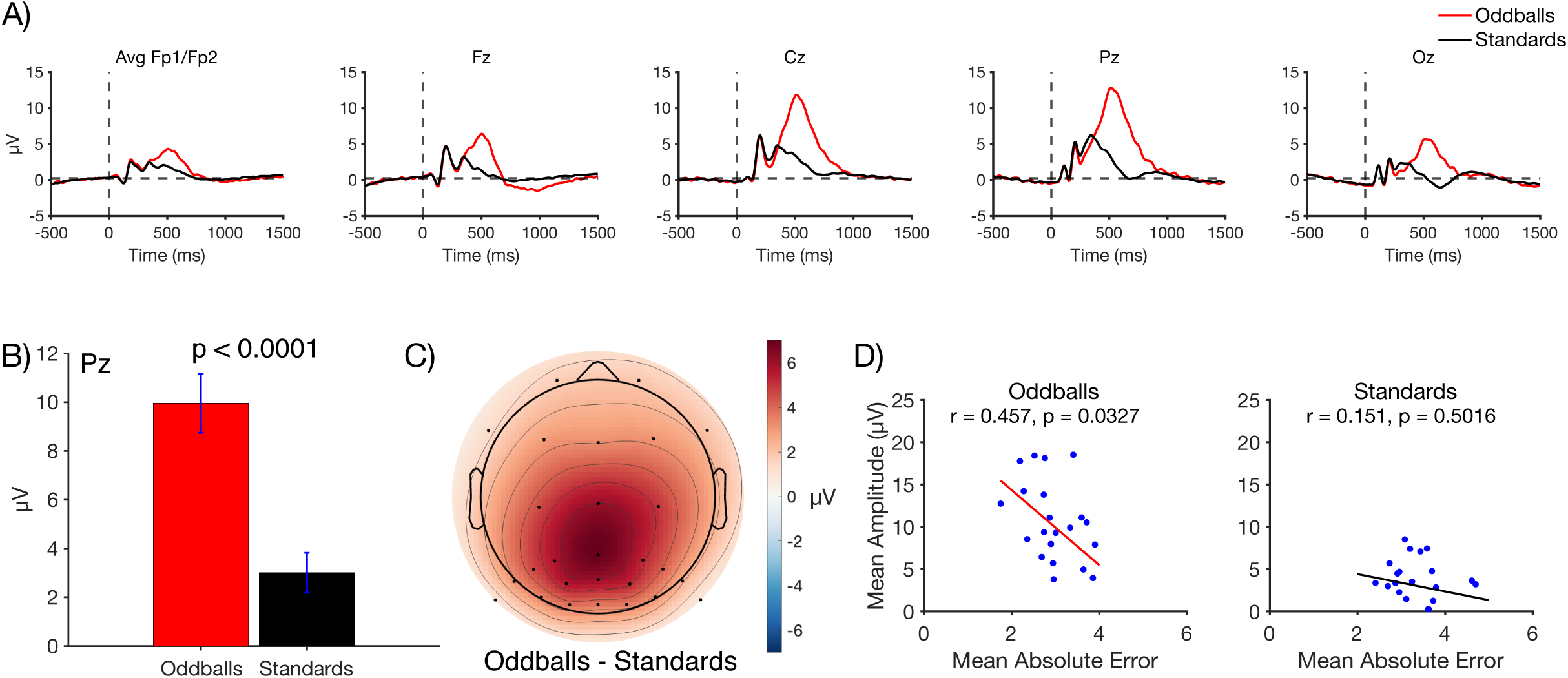
A) Grand average ERP waveforms for oddballs and standards at representative electrode sites. The P3b component peaked shortly after 500 ms and was largest at central and parietal electrode sites. B) Mean amplitude at site Pz measured across a 300-ms time window centered around the peak (362-662 ms). Error bars show ±1 SEM. C) Grand average scalp topography for the difference in amplitude between oddballs and standards in the P3b time window. D) Scatterplots showing the correlation between P3b amplitude (measured from 362-662 ms at the Pz electrode site) and behavioral accuracy on probe trials (mean absolute error). Each dot shows an individual participant, and the line is the best-fit regression line.

We also asked whether individuals with larger P3b amplitudes had more accurate working memory representations. Specifically, we examined the correlation between P3b amplitude (mean voltage at Pz during the measurement window on all oddball or standard trials) and the absolute response error (on oddball or standard probe trials). The scatterplots are shown in Figure 3d. A statistically significant correlation was observed for oddballs (*r* = 0.457, *p* = 0.033), with a smaller error for participants with a larger P3b amplitude. A correlation in the same direction was observed for standards, but it was weak and not significant (*r* = 0.151, *p* = 0.052).

We also asked whether trial-by-trial variations in P3b amplitude were related to behavioral performance on probe trials. To address this question, we performed a median split on single-trial P3b amplitudes to identify small-P3b and large-P3b trials for each participant, separately for standards and oddballs. We then computed the mean absolute error separately for the small-P3b trials and the large-P3b trials. As shown in Figure 4A, mean P3b amplitude was substantially smaller on small-P3b trials than on large-P3b trials, as expected given that these trials were sorted according to P3b amplitude. Figure 4B shows that behavioral performance was slightly worse on the small-P3b trials than on the large-P3b trials, but the difference was small and did not approach significance for either standards ((*t*(21) = 0.76, *p* = 0.46, *d_z_* =0.15) or oddballs (*t*(21) = 0.72, *p* = 0.71, *d_z_* =0.11)^2^.

**Figure 4.**
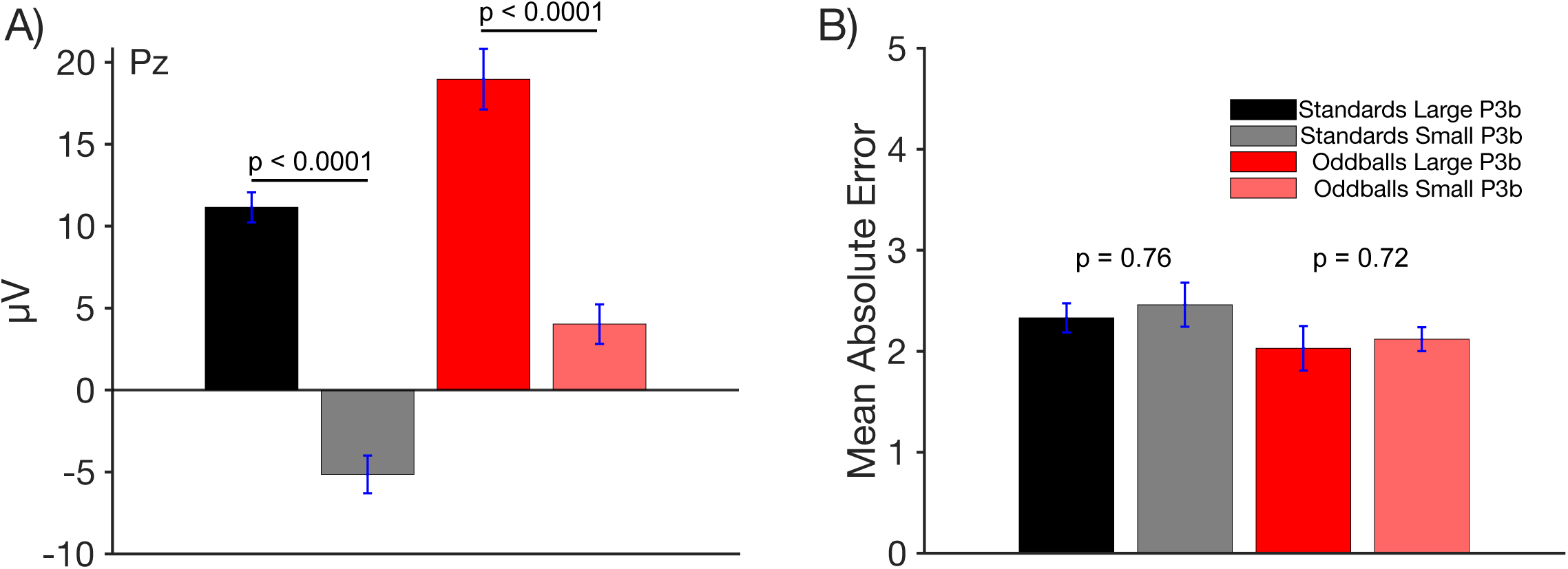
Median split of data based on P300 Amplitude. (A) Mean P3b amplitude for trials categorized as small-P3b and large-P3b, based on a median split of single-trial P3b amplitudes. As expected, small-P3b trials exhibited lower P3b amplitudes than large-P3b trials, with a significant difference observed for both oddballs and standards. (B) Mean absolute error in behavioral performance for small-P3b and large-P3b trials. Oddballs that produced larger P3b amplitudes were associated with slightly lower error rates, but this difference was not statistically significant. Error bars show ±1 SEM.

Together, these results show that oddball stimuli produce both a larger P3b and more accurate working memory representations, and that participants with larger P3b amplitudes for oddballs also have more accurate working memory responses. Note, however, that this does not demonstrate a causal relationship between the P3b and the working memory enhancement. It is entirely plausible that both effects are results of a common underlying factor (e.g., an increase in attention triggered by the oddballs, which separately impacts P3b amplitude and working memory encoding). Moreover, working memory performance was not significantly better on trials with large P3b amplitudes than on trials with small P3b amplitudes.

#### Decoding Results

Our final analyses were designed to determine whether we could see evidence of enhanced working memory for oddballs in the brain activity measured between the onset of the P3b wave and the end of the trial. Toward that end, we attempted to decode the location of the stimulus from the ERP activity at each moment in time across the recording epoch. We collapsed across the two locations that were near a given axis (e.g., 44° and 46°), giving us four cardinal locations and four diagonal locations. We then decoded which of the four cardinal locations was presented when the cardinals were standards, which of the four cardinal locations was presented when the cardinals were oddballs, which of the four diagonal locations was presented when the diagonals were standards, and which of the four diagonal locations was presented when the diagonal were oddballs. We then collapsed across the diagonal and cardinal categories to obtain overall decoding accuracy for oddballs and for standards. Because there were four locations within a category, chance decoding accuracy was 25%.

Figure 5a shows decoding accuracy at each individual time point. Location decoding accuracy was slightly greater for oddballs than for standards for much of the epoch, but most noticeably from approximately 800-1400 ms. To maximize statistical power, we averaged the decoding accuracy across an a priori time window that began at the start of the P3b measurement window (362 ms) and extended through the end of the epoch (1496 ms). As shown in Figure 5b, mean decoding accuracy was greater for oddballs than for standards, which was a significant difference in a paired *t* test (*t*(21) = 2.10, *p* = 0.048, *d_z_* = 0.425). Thus, the representation of the target was modestly but significantly enhanced for oddballs relative to standards during the period of time following stimulus offset.

**Figure 5.**
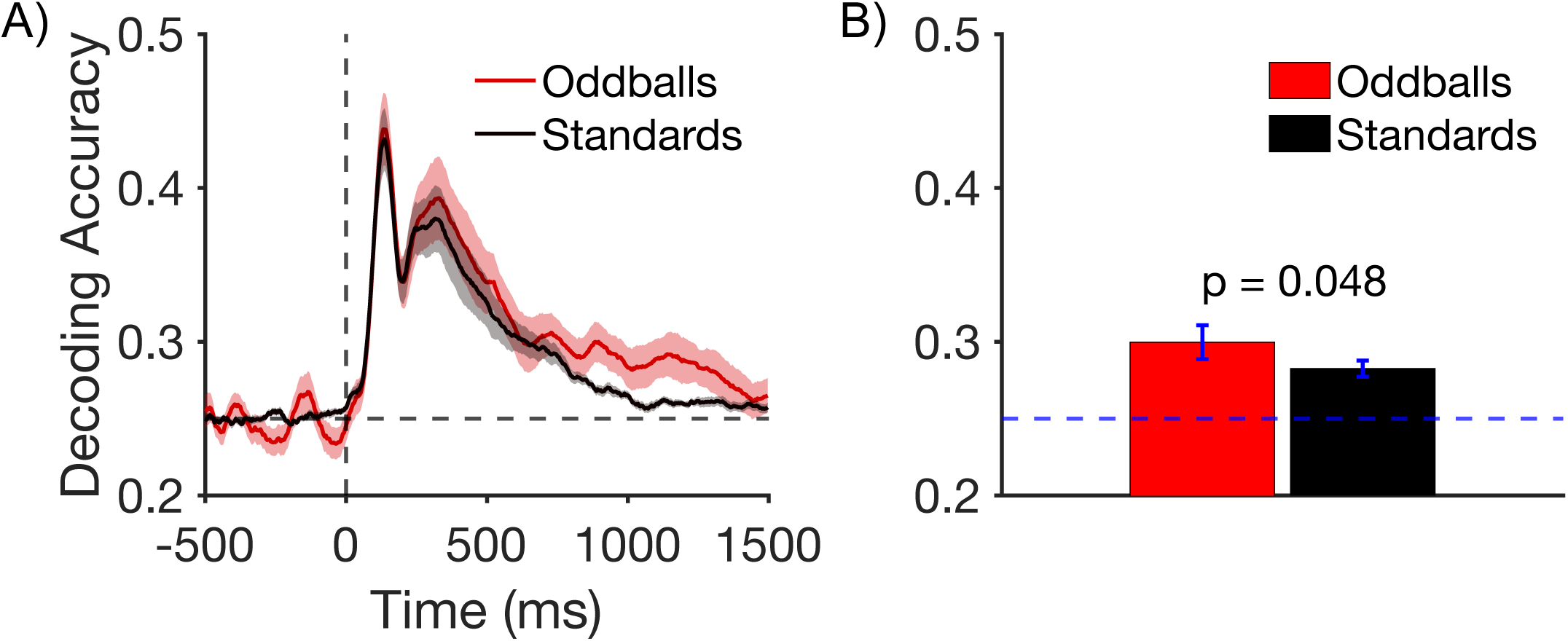
A) Timepoint-by-timepoint decoding of target location on oddball and standard trials. The dark lines show mean decoding accuracy across participants, and the shading shows ±1 SEM. B) Mean decoding accuracy between the start of the P3b measurement window (362 ms) and the end of the epoch (1500 ms). Error bars show ±1 SEM.

## Discussion

As expected, oddballs elicited a much larger P3b wave than standards, with the prototypical P3b scalp distribution. This indicates that the modifications we made to the oddball paradigm did not disrupt its fundamental nature. The behavioral data from probe trials showed that participants maintained a more accurate and less biased representation of the target for oddballs than for standards. In addition, the EEG decoding results showed that the neural representation of the target was enhanced for oddballs relative to standards in the period following P3b onset (although this effect was small). These findings are, to our knowledge, the first direct evidence that working memory is enhanced for rare, task-relevant stimulus categories.

ERP decoding methods are very sensitive, and it is important to consider the possibility that decoding accuracy in the present study was impacted by voltages other than those reflecting the neural activity underlying working memory. One such possibility would be electrooculographic voltages accompanying small but systematic eye movements in the direction of the target. We addressed this possibility by using ICA to correct ocular artifacts, which should have minimized the contribution of electrooculographic voltages to decoding accuracy. Another possibility would be neural activity related to preparing the manual response for reporting the location of the target on probe trials. This was addressed by the experimental design, which decreased the likelihood of significant preparation of motor responses in the direction of the target, because these responses were required on only 12.5% of trials (the probe trials). Moreover, probe responses occurred after the initial standard/oddball response, long after the time window in which we examined decoding, decreasing the likelihood that response preparation could have impacted decoding accuracy. Thus, it is unlikely that eye movements or response preparation contributed meaningfully to the decoding and could explain the difference in decoding accuracy between standards and oddballs. Moreover, even if these were the sources of the difference in decoding accuracy, the results would still indicate that the representation of the target location was enhanced for the oddball trials.

One possible alternative explanation for the difference in behavioral performance between the oddballs and the standards is proactive interference. Proactive interference occurs when the to-be-remembered target on the current trial is similar to targets from recent trials, leading to confusion at the time of recall (Baddeley, 1986; Keppel & Underwood, 1962). In the present study, locations in the standard category were, by design, presented more frequently than the locations in the oddball category, which could have led to more proactive interference and therefore worse performance for the standards than for the oddballs. This could not easily explain the difference in decoding accuracy between oddballs and standards, which occurred during the retention interval following the target, but it could explain the difference in behavioral probe accuracy.

## Experiment 2

To address the possibility of proactive interference in Experiment 1, we designed a new experiment in which some of the locations within the frequent category occurred with the same probability as the locations within the oddball category. Specifically, as illustrated in Figure 6A, we used visible boundary lines to trisect the circle into three regions. In a given block of trials, one of these regions was designated the oddball region and the other two were designated the standard region. Participants were instructed to press one button for oddballs and another button for standards. Stimuli appeared in the oddball region on 11.1% of trials. Unbeknownst to the participants, stimuli appeared in one of the two standard regions on 11.1% of trials (which we call *rare standards*) and in the other standard region on 77.8% of trials (which we call *frequent standards*). When debriefed at the end of the session, participants reported no awareness of the fact that one of the standard regions was rare and the other was frequent.

**Figure 6.**
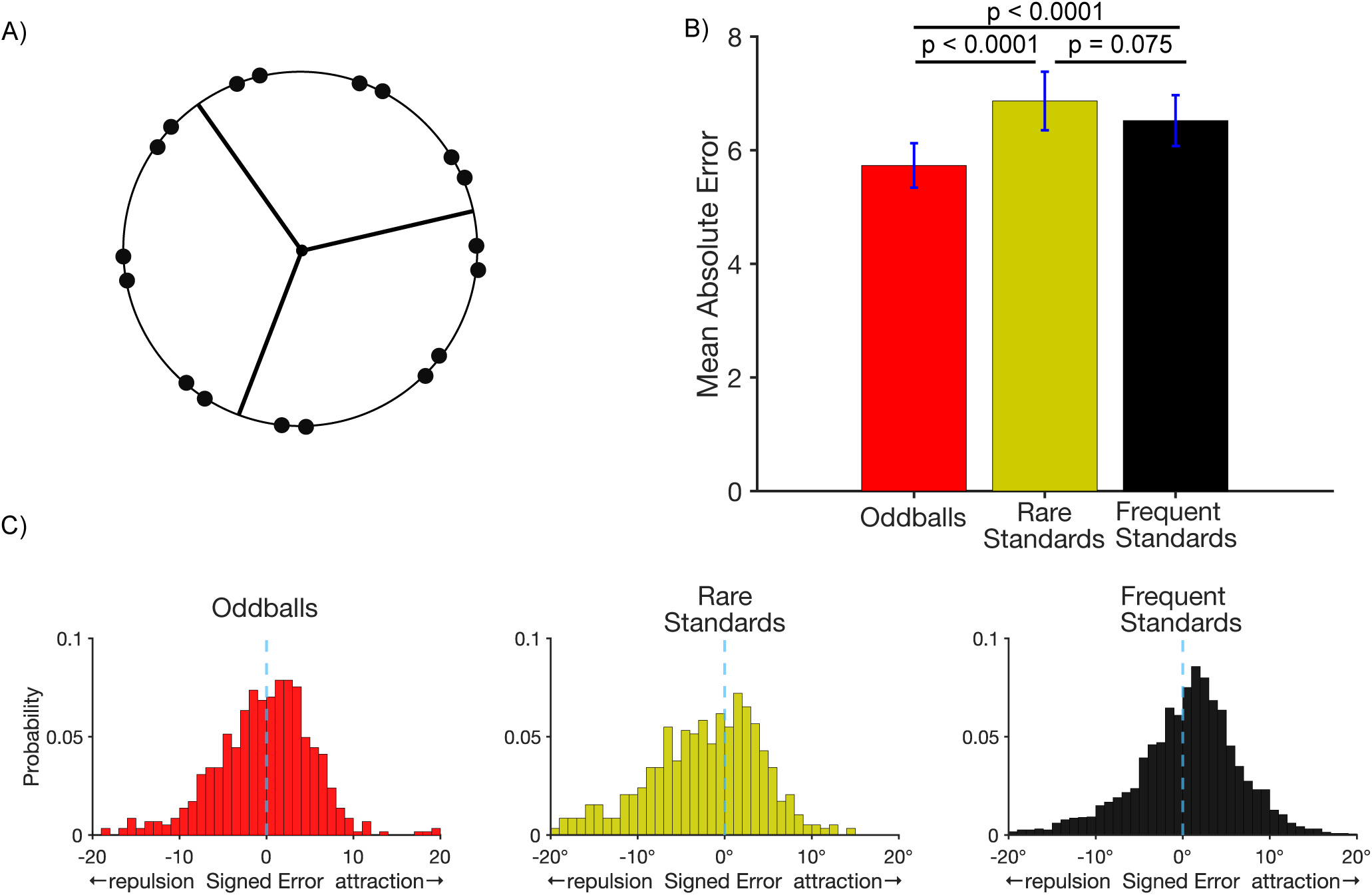
A) Stimulus setup for Experiment 2. On every trial, participants made a speeded keypress following the presentation of a target disc to indicate whether it appeared in one of 3 regions denoted by the lines. Targets appeared infrequently in two of the regions and frequently in the other. The frequent region and one of the rare regions shared the same button response frequent and rare standards. A different response was required for stimuli in the other rare region (oddballs). Each of these categories was followed by a probe at rate of 14.8%. B) Mean absolute error in the report of the exact location of the target disc on probe trials. Error bars show ±1 SEM. C) Histograms of the single-trial bias values for target discs near one of the boundary lines.

Although we did not record ERPs, it is well established that P3b amplitude reflects the probability of each task-defined category as a whole, not the probability of the specific stimuli (Luck, 2014; Luck & Hillyard, 1994; Mecklinger & Ullsperger, 1993), so it is reasonable to assume that the rare standard and frequent standards elicited approximately equivalent P3b amplitudes. However, because the rare oddball locations and rare standard locations occurred with equal probability, proactive interference should have been equivalent for the memories of the specific rare oddball and rare standard locations, which is what was assessed on the probe trials. If the enhanced behavioral probe performance for oddballs relative to standards in Experiment 1 was a result of greater proactive interference for the standards, then we should observe equivalent probe performance for the rare oddballs and the rare standards in the present experiment. If, however, rare task-defined categories lead to enhanced working memory relative to frequent task-defined categories, then we should observe enhanced probe performance for the rare oddballs than for the rare standards in the present experiment.

### Methods

#### Participants

Sixty human participants from UC Davis completed the study (49 women, and 11 men; mean age = 20.35, SD = 1.76). All participants were neurotypical and had normal or corrected-to-normal vision with no history of neurological conditions. 53 participants were right-handed, 6 were left-handed, and 1 was ambidextrous. Of these 60 participants, we excluded 3 due to accuracy being at chance for a given category, 5 due to experimenter errors, and 2 to a lack of probe trials following exclusion criteria). We chose to collect a larger sample of participants in this experiment than in Experiment 1 because of the smaller number of rare oddballs and rare standards in the present experiment. Compensation was provided via course credit. All participants provided informed consent, and the study was approved by the University of California, Davis Institutional Review Board.

#### Stimuli and Task

Figure 6a illustrates the task. Stimuli were presented using Psychopy (Peirce et al., 2019) with the same monitor, viewing distance, timing, response parameters and counterbalancing approaches as in Experiment 1. The disc appeared on the circle at locations separated by 40° (0°, 40°, 80°, 120°,160°, 200°, 240°, 280°, 320°) with ±1° of jitter (e.g., the stimulus was actually at either 39° or 41° for the location labeled 40°). Thus, there were 18 possible target locations. Note that orientations are quantified with respect to the unit circle, with 0° extending horizontally to the right from the fixation point.

Black boundary lines were continuously visible, extending from the fixation point to the perimeter of the circle at orientations of 100°, 220°, and 340°, which divided the 18 possible target locations into three regions. At the beginning of each trial block, one of the three regions was chosen as the oddball region, and participants were told to respond with one finger on each trial if the target appeared in that region and with another finger if the target appeared in either of the other two regions. Half the participants pressed the left arrow key for targets in the oddball region and the right arrow key for targets in the standard regions; this was reversed for the other half. The participants were told nothing about the probabilities of the locations in the different regions, and they were not exposed to terms such as “oddball,” “rare standard,” and “frequent standard.” They were simply told which button to press for a given region.

Each participant received 3 blocks of trials, and each region served as the oddball region in one block. The order of oddball regions was counterbalanced across participants. One of the other two regions was chosen to be the rare standard region and the other was the frequent standard region and switched off between blocks until all 3 regions had served as each trial type. Each block contained 324 trials: 36 targets in the oddball region, 36 targets in the rare standard region, and 252 targets in the frequent standard region. A random14.8% of trials were followed by a probe, resulting in a total of 18 oddball probe trials, 18 rare standard probe trials, and 108 frequent standard probe trials per participant.

#### Behavioral Analyses

For the oddball categorization task, we computed the proportion correct and the mean response time (RT) for each participant, separately for the oddball target, rare standard, and frequent standard categories.

For the probe task, the stimuli and responses were coded in terms of their angular position around the circle of possible target locations. As in experiment 1, we excluded trials for which the reported location was > 40° away from the true location (3.02% ± 1.45% of trials). We computed the mean absolute error for all positions and looked at the biases in the signed error for positions close to one of the boundaries.

### Results

#### Oddball task

We first examined the speed and accuracy for the button press task, in which participants indicated whether a given target appeared in the one region or in either of the other two regions. As expected given that the oddball response was much less frequent than the standard response, mean accuracy was significantly lower for rare oddballs (85.5% ± 1.30%) than for either rare standards (95.0% ± 0.71%; (*t*(49) = 8.40, *p* < 0.0001, *d_z_* = 1.28) or frequent standards (97.5% ± 0.79%; *t*(49) = 9.40, *p* < 0.0001, *d_z_* = 1.58). The small difference between rare standards and frequent standards was also statistically significant (*t*(49) = 5.15, *p* < 0.0001, *d_z_* = 0.47). In addition, mean RTs were significantly slower for rare oddballs (359.98 ± 11.68 ms) than for rare standards (304.75 ± 11.37 ms; *t*(49) = 9.82, *p* < 0.0001, *d_z_* = 0.68) or frequent standards (227.43 ± 8.92 ms; *t*(49) = 20.89, *p* < 0.0001, *d_z_* = 1.80). . The smaller difference between rare standards and frequent standards was also statistically significant (*t*(49) = 18.20, *p* < 0.0001, *d_z_* = 1.07). These results replicate the patter observed in Experiment 1, and also show that the rare and frequent standards were much more similar to each other than to the oddballs. Note also that the mean RTs were much faster in this experiment than in Experiment 1, presumably because the task of categorizing stimuli into the oddball and standard categories was much simpler in the present experiment.

#### Probe task

We next examined accuracy on the probe trials, for which participants clicked on the remembered location of the target. Figure 6b shows that the mean absolute response error for the probe task was smaller for oddballs (5.73 ± 0.390°) than for either rare standards (6.87 ± 0.516°; *t*(49) = 3.85, *p* < 0.0001, *d_z_* = 0.35) or frequent standards (6.52 ± 0.448°; *t*(49) = 3.91, *p* < 0.0001, *d_z_* = 0.27)). The same pattern was found when the standard deviation of response errors was used as the metric of memory precision (supplementary Figure S1). The better performance for the oddballs than for the frequent standards replicates the pattern observed in Experiment 1, and the better performance for oddballs than for the rare standards demonstrates that oddballs yield enhanced memory performance even when the probability of the individual target locations was equivalent, thus controlling for proactive interference. There was no significant difference in probe response error between rare and frequent standards (*t*(49) = 1.82, *p* = 0.075, *d_z_* = 0.1).

As in Experiment 1, we asked whether working memory representations of oddballs are less subject to systematic biases than the rare and frequent standards. Whereas Experiment 1 examined this using the “virtual boundaries” of the horizontal and vertical axes, this was examined in Experiment 2 using the visible boundaries that trisected the circle. Within each of these regions, we examined biases for the pairs of locations that were close to the boundaries, excluding the pairs of locations that were in the center of each region. We coded each response error with a negative sign if the reported location was shifted away from the true location in a direction opposite to the nearest boundary and with a positive sign if the reported location was shifted toward the nearest boundary. As shown in Figure 2c, the response errors for all three types of targets exhibited a peak that was slightly in the direction of attraction toward the boundary but also exhibited a skew away from the boundary. For each participant, we computed the mean of these signed error values as an overall metric of bias.

For oddballs, the mean bias score was slightly skewed away from the boundary, but this effect was not significantly different from zero (*t*(49) = -2.00, *p* = 0.052, *d_z_* = -0.28). For rare standards, the distribution was more highly skewed away from the boundary, and the mean bias score was significantly different from zero (*t*(49) = -6.22, *p* < 0.0001, *d_z_* = -0.88). The distribution was not highly skewed for frequent standards, and the mean bias was not significantly different from zero (*t*(49) = -0.944, *p* = 0.350, *d_z_* = -0.13). The key finding was that the mean bias score was significantly more negative for rare standards than for oddballs (*t*(49) = 6.24, *p* < 0.0001, *d_z_* = 0.54). This demonstrates that the difference in reduced bias for oddballs relative to standards observed in Experiment 1 can also be observed when proactive interference is controlled by equating the probabilities of the individual locations. Curiously, the mean bias score was also significantly more negative for rare standards than for frequent standards (*t*(49) = 6.24, *p* < 0.0001, *d_z_* = 0.54), and there was a significant difference between oddballs and frequent standards (*t*(49) = 2.98, *p* = 0.004, *d_z_* = 0.17).

## Discussion

Experiment 2 was designed to assess an alternative explanation of the greater behavioral probe performance for oddballs than for standards in Experiment 1, namely a difference in proactive interference from the previous trials (which were more likely to be standards than to be oddballs). We addressed this in experiment 2 by comparing oddballs with a subset of standards that occurred with the same probability as the oddballs. The results from the probe trials indicate that, as in Experiment 1, participants maintained a more accurate working memory representation for locations in the oddball category than for locations in the standard category, even for the subset of standards that occurred with the same probability as the oddballs. This rules out differential proactive interference as an explanation for the enhanced memory performance for the oddballs.

The oddballs also led to less bias than the rare standards, mirroring the finding of less bias for oddballs than for standards in Experiment 1. Surprisingly, there was a difference in bias between the oddballs and the frequent standards. We have no explanation for this. However, the difference in bias between the oddballs and the rare standards indicates that items belonging to a rare task-defined category are remembered with less bias than equally rare items belonging to a frequent task-defined category. Note that the bias may have reflected repulsion away from the boundary line that separated the three regions or an attraction toward the center of the region (Gold et al., 2020; Huttenlocher et al., 1994; Schmidt et al., 2003); in either case, the bias was reduced for the oddballs relative to the rare standards. More generally, behavioral biases such as those observed here and in Experiment 1 may arise at many different stages of processing; no matter when they arise, they appear to be reduced for oddballs.

### General Discussion

The current study sought to provide direct evidence that working memory representations are enhanced for relatively rare task-relevant event categories compared to relatively frequent task-relevant event categories. Traditional oddball paradigms do not provide a sensitive assessment of working memory representations, but we were able to assess the accuracy of target representations behaviorally by including probe trials. We also assessed the neural representation of the target by decoding the target’s location on the basis of the EEG activity following target offset.

In Experiment 1, we observed a canonical P3b effect, with a greater amplitude for the rare category than for the frequent category at the Pz electrode site, with the prototypical P3b scalp distribution. This indicates that our oddball task, which was involved a relatively complex categorization task, yielded the same basic pattern of ERP effects as more typical oddball tasks. However, it is important to note that the added demand of needing to report the exact target locations on probe trials may have influenced the memory representations of the rare events, potentially enhancing the P3b response in ways that may differ from more standard oddball paradigms

The behavioral results from the probe trials provided a sensitive assay of the memory representations for the individual locations within the oddball and standard categories. We found that memory reports were more precise and less biased for locations within the oddball category than for locations within the standard category. Moreover, we found that the mean memory errors when oddball trials were probed were smaller in individuals with larger P3b amplitudes for the oddball stimuli. This provides a somewhat more direct link between the P3b and working memory performance (although the nature of this link is subject to all the limitations that arise from simple correlations). Finally, we saw in our decoding analyses that the neural representation of the oddball stimulus was enhanced following the onset of the P3b (albeit with a relatively small effect size). This is important, because it shows a difference in the representation during the delay period following the stimulus, whereas the behavioral findings could reflect processes that happen at the time of report. These findings are, to our knowledge, the first direct evidence that working memory is enhanced for stimuli that belong to rare, task-defined stimulus categories.

However, when we broke down the trials within the rare and frequent categories on the basis of P3b amplitude, we did not find enhanced working memory performance for trials with a larger P3b than for trials with a smaller P3b within either the rare category or the frequent category. In addition, whereas the P3b peaked at approximately 500 ms, the effects of rareness on decoding accuracy were largely evident after approximately 800 ms. This suggests that rareness impacts the maintenance of information in working memory, not the initial encoding.

Experiment 2 ruled out alternative explanations for the behavioral results, showing that behavioral performance was more accurate and less biased for stimuli within the oddball category than for equally probable stimuli within the standard category. Thus, it is the probability of the category as a whole, not the probability of the individual instances within a category, that determines the quality of the working memory representation.

The present results do not indicate the nature of the neural mechanism underlying enhanced working memory, but such an effect could potentially be a result of increased firing by neurons in the locus coeruleus, which would lead to a phasic increase in norepinephrine release throughout the cortex (Aston-Jones & Cohen, 2005; Krebs et al., 2018; Murphy et al., 2011; Nieuwenhuis et al., 2005). This is consistent with prior findings that norepinephrine is critical for signaling oddballs in humans and that norepinephrine depletion leads to working memory deficits in nonhuman primates (Arnsten, 2006; Arnsten & Goldman-Rakic, 1985; Brozoski et al., 1979; Cai et al., 1993; Franowicz et al., 2002; Strange & Dolan, 2007; Zhang et al., 2013). More generally, the enhanced working memory we observed for oddballs could reflect greater allocation of attention to rare stimuli. Indeed, suppression of alpha-band EEG activity, a well-known correlate of attentional allocation (Adrian & Matthews, 1934; Jensen & Mazaheri, 2010) is greater for oddballs than for standards (Klimesch et al., 1998; Yordanova et al., 2001).

The finding that working memory representations were enhanced for rare items—with greater accuracy for participants with larger P3b amplitudes—might be interpreted as evidence for the *context updating* hypothesis of the P3b wave, which was proposed over 40 years ago by Donchin (1981). Although Donchin did not use the term *working memory*, more recent researchers have interpreted the context updating hypothesis as stating that the P3b is related to working memory encoding (Kok, 2001; Polich, 2007). However, there is not much evidence to support the idea that the P3b specifically reflects the updating of working memory (Verleger, 2008). Indeed, using a *reference-back* task that was specifically designed to distinguish between updating and other processes, Rac-Lubashevsky and Kessler (2019) found no evidence that a P3b was triggered when working memory updating was required.

Moreover, the P3b component in the present study was more than three times larger for oddballs than for standards, and yet behavioral accuracy on probe trials (a fairly direct measure of working memory) was only slightly greater for oddballs than for standards. In other words, even though standards produce only a small P3b, there were encoded into working memory quite accurately. We also found no significant difference in working memory performance between trials with a large P3b and trials with a small P3b within either the standard category or the oddball category. In addition, although decoding accuracy greater for oddballs than for standards, the difference in decoding accuracy was modest and it was not evident until long after the onset of P3b activity.

Thus, it is unlikely that the neural mechanisms that produce the P3b component are the same mechanisms that encode information into working memory. The correlation between P3b amplitude and working memory accuracy observed in Experiment 1 is more likely related to a shared impact of attentional allocation on P3b amplitude and working memory encoding. By analogy, pupil dilation is also enhanced for oddballs, correlated with working memory performance, and linked with locus coeruleus activity (Aminihajibashi et al., 2019; Eckstein et al., 2019; Gilzenrat et al., 2010; Murphy et al., 2011; Robison et al., 2023; Robison & Unsworth, 2019; Zokaei et al., 2019). However, this does not mean that the mechanisms that produce pupil dilation are the same as the mechanisms that encode information into working memory. Instead, the P3b may be related to models in which locus coeruleus activity provides a signal across the cerebral cortex that predictions about the environment have been strongly violated (Jordan, 2023). Pupil dilation has been linked to the P3b ERP component, with a positive correlation between increases in the P3b amplitude and task evoked pupil dilation (Contier et al., 2024), but additional research would be needed to provide conclusive evidence of causal links between locus coeruleus activity and the P3b wave.

Behavioral responses in working memory tasks can be influenced by decision processes at the time of report in addition to encoding and maintenance processes. To provide additional information about the nature of the differences in working memory between rare and frequent stimuli, we decoded the location of the target being held in working memory using the scalp topography of the ERP signal during the interval immediately following the target, prior to the probe period. The finding of greater decoding accuracy for oddballs than for standards during this period suggests that rare stimuli are *encoded* and *maintained* more accurately than frequent stimuli (as opposed to an effect of rareness that is isolated to decision processes at the time of response). However, we cannot be certain that these ERP decoding effects reflect the same processes responsible for the improved behavioral performance for rare stimuli on probe trials. Indeed, the ERP decoding analyses were designed to distinguish among the four different locations within a given response category, which were separated by 90°, whereas the behavioral responses were typically within 5° of the true target location (see Figure 2A). The coarseness of the decoding process may partly explain why the differences in decoding accuracy between oddballs and standards were so small. We are not yet at the point where we can decode differences in stimulus location from scalp voltages with the same precision as the behavioral responses.

We would like to emphasize that the present study used a behavioral task to create the rare and frequent stimulus categories. These were not categories that vary in their frequency or novelty outside of this task. Stimuli that are intrinsically novel elicit a very different pattern of neural activity, including the novelty P3a component (which has a much more frontal scalp distribution than the P3b component) and extensive BOLD activity in temporal and inferior frontal cortex (Strobel et al., 2008). It is not known how intrinsic novelty impacts working memory. Performance in visual working memory tasks is more accurate for familiar than for unfamiliar stimuli under some conditions (Chen et al., 2006; Jackson & Raymond, 2008; Ngiam et al., 2019), but this likely reflects the use of longer-term memory representations to aid in task performance. Novelty can improve working memory under other conditions, particularly encoding processes (Mayer et al., 2011). Additional research will be necessary to fully understand how this kind of intrinsic novelty is related to the encoding and maintenance of information leading to enhancements in working memories.

In addition, different motor responses were made for the rare and frequent stimulus categories in the present experimental design, and it is possible that the rareness of the motor response rather than the rareness of the stimulus category was responsible for the difference in working memory performance and decoding accuracy in the present study. Indeed, substantial evidence indicates that, although the P3b does not require a motor response (Verleger et al., 2016), response-related processes have a large impact on the P3b component (Verleger, 2020). Additional research will be needed to determine whether the differences in working memory between standards and oddballs in the present study were a result of differences in the probability of the motor responses.

## Data and Code Availability

Presentation scripts, EEG/ERP data, behavioral data and analysis scripts used are available at doi.org/10.17605/OSF.IO/HV7JU.

## Author Contributions

Carlos Daniel Carrasco: conceptualization, experimental design, data collection, analyses, writing; Aaron Mathew Simmons: software development, data collection; John E. Kiat: analyses; Steven J. Luck: conceptualization, experimental design, writing, funding acquisition. Editing/Review was performed by all authors prior to submission of the final version of the manuscript.

## Footnotes

1 Note that considerable evidence suggests that the temporal probability rather than the sequential probability is the key determinant of P3b amplitude (Gonsalvez et al., 2007; Gonsalvez & Polich, 2002). These two types of probability covary in the present study (and most oddball experiments), and it does not matter for the present study whether the key variable is sequential or temporal probability.

2 This median-split analysis dichotomizes P3b amplitude into categories of small and large, and this approach may have less statistical power than treating P3b amplitude as a continuous variable. We therefore also performed a linear mixed effects analysis on the single-trial data to ask whether P3b amplitude was correlated with the absolute error of the behavioral response. We again found no significant effect.

## Notes

The authors declare no competing financial interests.

https://www.doi.org/10.17605/OSF.IO/HV7JU

### Competing Interest Statement

The authors have declared no competing interest.

### Summary of Updates

Revision based on feedback from reviewers prior to submission for publication in psychophysiology.

## References

Adrian, E. D., & Matthews, B. H. C. (1934). The Berger rhythm: potential changes from the occipital lobes in man. Brain, 57(4), 355–385.

Aminihajibashi, S., Hagen, T., Foldal, M. D., Laeng, B., & Espeseth, T. (2019). Individual differences in resting-state pupil size: Evidence for association between working memory capacity and pupil size variability. International Journal of Psychophysiology, 140, 1–7. 10.1016/j.ijpsycho.2019.03.007

Arnsten, A. F. T. (2006). Fundamentals of Attention-Deficit/Hyperactivity Disorder: Circuits and Pathways. In J Clin Psychiatry (Vol. 67, Issue 8).

Arnsten, A. F. T., & Goldman-Rakic, P. S. (1985). α2-Adrenergic Mechanisms in Prefrontal Cortex Associated with Cognitive Decline in Aged Nonhuman Primates. In New Series (Vol. 13, Issue 4731).

Aston-Jones, G., & Cohen, J. D. (2005). An integrative theory of locus coeruleus-norepinephrine function: Adaptive gain and optimal performance. In Annual Review of Neuroscience (Vol. 28, pp. 403–450). 10.1146/annurev.neuro.28.061604.135709

Baddeley, A. (1986). Working memory. Oxford: Oxford University Press, Clarendon Press.

Bae, G. Y. (2022). Breaking the cardinal rule: The impact of interitem interaction and attentional priority on the cardinal biases in orientation working memory. *Attention*, Perception, and Psychophysics, 84(7), 2186–2194. 10.3758/s13414-021-02374-2

Bae, G. Y., & Luck, S. J. (2019). Decoding motion direction using the topography of sustained ERPs and alpha oscillations. NeuroImage, 184, 242–255. 10.1016/j.neuroimage.2018.09.029

Bae, G. Y. (2021). The Time Course of Face Representations during Perception and Working Memory Maintenance. Cerebral Cortex Communications, 2(1), 1–12. 10.1093/texcom/tgaa093

Bae, G. Y., & Luck, S. J. (2018). Dissociable Decoding of Spatial Attention and Working Memory from EEG Oscillations and Sustained Potentials. The Journal of Neuroscience, 38(2), 2860–17. 10.1523/JNEUROSCI.2860-17.2017

Bledowski, C., Prvulovic, D., Goebel, R., Zanella, F. E., & Linden, D. E. J. (2004). Attentional systems in target and distractor processing: A combined ERP and fMRI study. NeuroImage, 22(2), 530–540. 10.1016/j.neuroimage.2003.12.034

Brozoski, T. J., Brown, R. M., Rosvold, H. E., & Goldman, P. S. (1979). Cognitive Deficit Caused by Regional Depletion of Dopamine in Prefrontal Cortex of Rhesus Monkey. In New Series (Vol. 205, Issue 4409).

Cai, J. X., Ma, Y.-Y., & Hu, X.-T. (1993). Reserpine impairs spatial working memory performance in monkeys: reversal by the az-adrenergic agonist clonidine. In Brain Research (Vol. 614).

Chen, D., Yee Eng, H., & Jiang, Y. (2006). Visual working memory for trained and novel polygons. Visual Cognition, 14(1), 37–54. 10.1080/13506280544000282

Clark, V. P., Fannon, S., Lai, S., Benson, R., & Bauer, L. (2000). Responses to Rare Visual Target and Distractor Stimuli Using Event-Related fMRI.

Cohen J. (1988). Statistical Power Analysis for the Behavioral Sciences Second Edition.

Contier, F., Wartenburger, I., Weymar, M., & Rabovsky, M. (2024). Are the P600 and P3 ERP components linked to the task-evoked pupillary response as a correlate of norepinephrine activity? Psychophysiology, 61(7). 10.1111/psyp.14565

Delorme, A., & Makeig, S. (2004). EEGLAB: an open source toolbox for analysis of single-trial EEG dynamics including independent component analysis. In Journal of Neuroscience Methods (Vol. 134). http://www.sccn.ucsd.edu/eeglab/

Dietterich, T. G., & Bakiri, G. (1995). Solving Multiclass Learning Problems via Error-Correcting Output Codes. In Journal of Artiicial Intelligence Research (Vol. 2).

Donchin, E. (1981). Surprise!…Surprise? Psychophysiology, 18(5), 493–513. 10.1111/j.1469-8986.1981.tb01815.x

Donchin, E., & Coles, M. G. H. (1988). Is the P300 component a manifestation of context updating? Behavioral and Brain Sciences, 11(3), 357–374. 10.1017/S0140525X00058027

Eckstein, M. K., Starr, A., & Bunge, S. A. (2019). How the inference of hierarchical rules unfolds over time. Cognition, 185, 151–162. 10.1016/j.cognition.2019.01.009

Franowicz, J. S., Kessler, L. E., Dailey Borja, C. M., Kobilka, B. K., Limbird, L. E., & Arnsten, A. F. T. (2002). Mutation of the 2A-Adrenoceptor Impairs Working Memory Performance and Annuls Cognitive Enhancement by Guanfacine.

Gilzenrat, M. S., Nieuwenhuis, S., Jepma, M., & Cohen, J. D. (2010). Pupil diameter tracks changes in control state predicted by the adaptive gain theory of locus coeruleus function. *Cognitive*, Affective and Behavioral Neuroscience, 10(2), 252–269. 10.3758/CABN.10.2.252

Gold, J. M., Bansal, S., Anticevic, A., Cho, Y. T., Repovš, G., Murray, J. D., Hahn, B., Robinson, B. M., & Luck, S. J. (2020). Refining the empirical constraints on computational models of spatial working memory in schizophrenia. Biological Psychiatry: Cognitive Neuroscience and Neuroimaging, 5(9), 913–922.

Gonsalvez, C. J., Barry, R. J., Rushby, J. A., & Polich, J. (2007). Target-to-target interval, intensity, and P300 from an auditory single-stimulus task. Psychophysiology, 44(2), 245–250.

Gonsalvez, C. J., & Polich, J. (2002). P300 amplitude is determined by target-to-target interval. Psychophysiology, 39(3), 388–396.

Huttenlocher, J., Newcombe, N., & Sandberg, E. H. (1994). The coding of spatial location in young children. Cognitive Psychology, 27(2), 115–147.

Jackson, M. C., & Raymond, J. E. (2008). Familiarity Enhances Visual Working Memory for Faces. Journal of Experimental Psychology: Human Perception and Performance, 34(3), 556–568. 10.1037/0096-1523.34.3.556

Jensen, O., & Mazaheri, A. (2010). Shaping functional architecture by oscillatory alpha activity: Gating by inhibition. Frontiers in Human Neuroscience, 4. 10.3389/fnhum.2010.00186

Johnson, R. (1988). SCALP-RECORDED P300 ACTIVITY IN PATIENTS FOLLOWING UNILATERAL TEMPORAL LOBECTOMY. In Brain (Vol. 111). https://academic.oup.com/brain/article/111/6/1517/297108

Jordan, R. (2023). The locus coeruleus as a global model failure system. In Trends in Neurosciences. Elsevier Ltd. 10.1016/j.tins.2023.11.006

Katayama, J., & Polich, J. (1998). Stimulus context determines P3a and P3b. Psychophysiology, 35(1), 23–33. 10.1111/1469-8986.3510023

Keppel, G., & Underwood, B. J. (1962). Proactive Inhibition in Short-Term Retention of Single Items 1.

Kim, H. (2014). Involvement of the dorsal and ventral attention networks in oddball stimulus processing: A meta-analysis. Human Brain Mapping, 35(5), 2265–2284. 10.1002/hbm.22326

Klimesch, W., Doppelmayr, M., Russegger, H., Pachinger, T., & Schwaiger, J. (1998). Induced alpha band power changes in the human EEG and attention. Neuroscience Letters, 244(2), 73–76.

Kok, A. (2001). On the utility of P3 amplitude as a measure of processing capacity. Psychophysiology, 38(3), 557–577. 10.1017/S0048577201990559

Krebs, R. M., Park, H. R. P., Bombeke, K., & Boehler, C. N. (2018). Modulation of locus coeruleus activity by novel oddball stimuli. Brain Imaging and Behavior, 12(2), 577–584. 10.1007/s11682-017-9700-4

Kutas, M., Mccarthy, G., & Donchin, E. (1977). Augmenting Mental Chronometry: The P300 as a Measure of Stimulus Evaluation Time. In Source: Science (Vol. 197, Issue 4305).

Lakens, D. (2013). Calculating and reporting effect sizes to facilitate cumulative science: A practical primer for t-tests and ANOVAs. Frontiers in Psychology, 4(NOV). 10.3389/fpsyg.2013.00863

Linden, D. E. J., Prvulovic, D., Formisano, E., Völlinger, M., Zanella, F. E., Goebel, R., & Dierks, T. (1999). The functional neuroanatomy of target detection: an fMRI study of visual and auditory oddball tasks. Cerebral Cortex, 9(8), 815–823.

Lopez-Calderon, J., & Luck, S. J. (2014). ERPLAB: An open-source toolbox for the analysis of event-related potentials. Frontiers in Human Neuroscience, 8(1 APR), 1–14. 10.3389/fnhum.2014.00213

Luck, S. J. (2014). An Introduction To The Event-Related Potential Technique (2nd Editio). MIT Press Journals.

Luck, S. J. (2023). ERPLAB Decoding Tutorial. https://github.com/ucdavis/erplab/wiki/ERPLAB-Decoding-Tutorial

Luck, S. J., & Hillyard, S. A. (1994). Electrophysiological correlates of feature analysis during visual search. Psychophysiology, 31(3), 291–308.

Mayer, J. S., Kim, J., & Park, S. (2011). Enhancing visual working memory encoding: The role of target novelty. Visual Cognition, 19(7), 863–885. 10.1080/13506285.2011.594459

Mecklinger, A., & Ullsperger, P. (1993). P3 varies with stimulus categorization rather than probability. In Electroencephalography and clinical Neurophysiology (Vol. 86).

Murphy, P. R., Robertson, I. H., Balsters, J. H., & O’connell, R. G. (2011). Pupillometry and P3 index the locus coeruleus-noradrenergic arousal function in humans. Psychophysiology, 48(11), 1532–1543. 10.1111/j.1469-8986.2011.01226.x

Ngiam, W. X. Q., Khaw, K. L. C., Holcombe, A. O., & Goodbourn, P. T. (2019). Visual working memory for letters varies with familiarity but not complexity. Journal of Experimental Psychology: Learning, Memory, and Cognition, 45(10), 1761.

Nieuwenhuis, S., Aston-Jones, G., & Cohen, J. D. (2005). Decision making, the P3, and the locus coeruleus-norepinephrine system. In Psychological Bulletin (Vol. 131, Issue 4, pp. 510–532). 10.1037/0033-2909.131.4.510

Nieuwenhuis, S., De Geus, E. J., & Aston-Jones, G. (2011). The anatomical and functional relationship between the P3 and autonomic components of the orienting response. In Psychophysiology (Vol. 48, Issue 2, pp. 162–175). Blackwell Publishing Inc. 10.1111/j.1469-8986.2010.01057.x

Paller, K. A., Mccarthy, G., Roessler, E., Allison, T., & Wood, C. C. (1992). Potentials evoked in human and monkey medial temporal lobe during auditory and visual oddball paradigms. In Electroencephalography and clinical Neurophysiology (Vol. 84).

Peirce, J., Gray, J. R., Simpson, S., MacAskill, M., Höchenberger, R., Sogo, H., Kastman, E., & Lindeløv, J. K. (2019). PsychoPy2: Experiments in behavior made easy. Behavior Research Methods, 51(1), 195–203. 10.3758/s13428-018-01193-y

Polich, J. (1986). A’ITENTION, PROBABILITY, AND TASK DEMANDS AS DETERMINANTS OF P300 LATENCY FROM AUDITORY STIMULI t. In Electroencephalograph)’ and clinical Neurophysiology (Vol. 63).

Polich, J. (2007). Updating P300: An integrative theory of P3a and P3b. Clinical Neurophysiology, 118(10), 2128–2148. 10.1016/j.clinph.2007.04.019

Polich, J. (2012). Neuropsychology of P300. The Oxford Handbook of Event-Related Potential Components, 159–188.

Pratte, M. S., Park, Y. E., Rademaker, R. L., & Tong, F. (2017). Accounting for stimulus-specific variation in precision reveals a discrete capacity limit in visual working memory. Journal of Experimental Psychology: Human Perception and Performance, 43(1), 6–17. 10.1037/xhp0000302

Rac-Lubashevsky, R., & Kessler, Y. (2019). Revisiting the relationship between the P3b and working memory updating. Biological Psychology, 148. 10.1016/j.biopsycho.2019.107769

Robison, M. K., Ralph, K. J., Gondoli, D. M., Torres, A., Campbell, S., Brewer, G. A., & Gibson, B. S. (2023). Testing locus coeruleus-norepinephrine accounts of working memory, attention control, and fluid intelligence. *Cognitive*, Affective and Behavioral Neuroscience, 23(4), 1014–1058. 10.3758/s13415-023-01096-2

Robison, M. K., & Unsworth, N. (2019). Pupillometry tracks fluctuations in working memory performance. *Attention*, Perception, and Psychophysics, 81(2), 407–419. 10.3758/s13414-018-1618-4

Schmidt, T., Werner, S., & Diedrichsen, J. (2003). Spatial distortions induced by multiple visual landmarks: How local distortions combine to produce complex distortion patterns. Perception & Psychophysics, 65(6), 861–873.

Soltani, M., & Knight, R. T. (2000). Neural Origins of the P300. In Critical Reviews in Neurobiology (Vol. 14).

Strange, B. A., & Dolan, R. J. (2007). β-adrenergic modulation of oddball responses in humans. Behavioral and Brain Functions, 3. 10.1186/1744-9081-3-29

Strobel, A., Debener, S., Sorger, B., Peters, J. C., Kranczioch, C., Hoechstetter, K., Engel, A. K., Brocke, B., & Goebel, R. (2008). Novelty and target processing during an auditory novelty oddball: A simultaneous event-related potential and functional magnetic resonance imaging study. NeuroImage, 40(2), 869–883. 10.1016/j.neuroimage.2007.10.065

Verleger, R. (2008). P3b: Towards some decision about memory. In Clinical Neurophysiology (Vol. 119, Issue 4, pp. 968–970). 10.1016/j.clinph.2007.11.175

Verleger, R. (2020). Effects of relevance and response frequency on P3b amplitudes: Review of findings and comparison of hypotheses about the process reflected by P3b. Psychophysiology, 57(7). 10.1111/psyp.13542

Verleger, R., Grauhan, N., & Śmigasiewicz, K. (2016). Go and no-go P3 with rare and frequent stimuli in oddball tasks: A study comparing key-pressing with counting. International Journal of Psychophysiology, 110, 128–136.

Wei, X. X., & Stocker, A. A. (2015). A Bayesian observer model constrained by efficient coding can explain “anti-Bayesian” percepts. Nature Neuroscience, 18(10), 1509–1517. 10.1038/nn.4105

Yordanova, J., Kolev, V., & Polich, J. (2001). P300 and alpha event-related desynchronization (ERD). Psychophysiology, 38(1), 143–152.

Zhang, Z., Cordeiro Matos, S., Jego, S., Adamantidis, A., & Séguéla, P. (2013). Norepinephrine Drives Persistent Activity in Prefrontal Cortex via Synergistic α1 and α2 Adrenoceptors. PLoS ONE, 8(6). 10.1371/journal.pone.0066122

Zokaei, N., Board, A. G., Manohar, S. G., & Nobre, A. C. (2019). Modulation of the pupillary response by the content of visual working memory. Proceedings of the National Academy of Sciences of the United States of America, 116(45), 22802–22810. 10.1073/pnas.1909959116

