## Supplementary Figure S1 for "Enhanced Working Memory Representations for Rare Events"

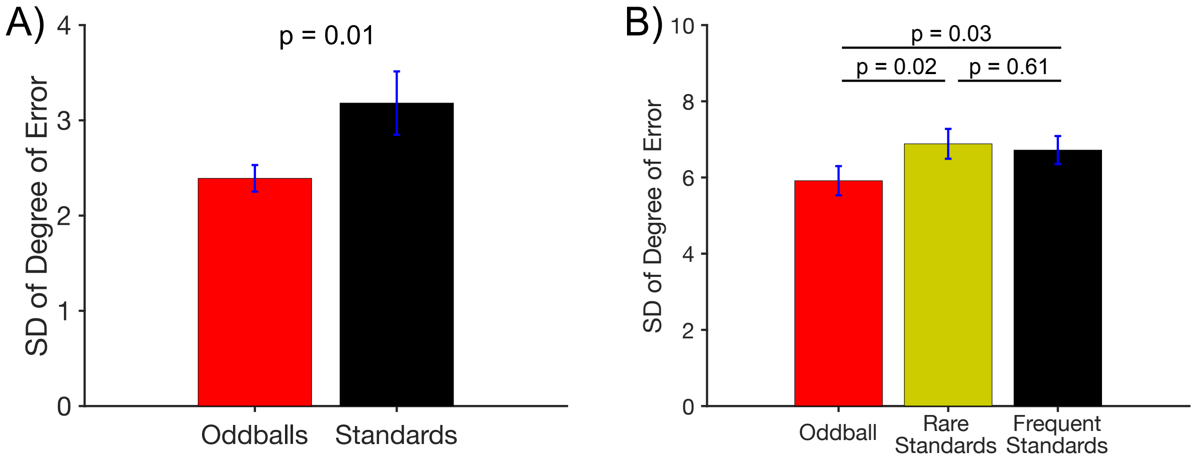


Figure S1. Additional metrics of working memory precision. A) Standard deviation (SD) of the response error in Experiment 1. As was observed for the mean absolute error, the SD was significantly greater for oddballs than standards $t(21) = 2.88$, $p = 0.01$, $d_z = 0.66$. B) SD of the response error in Experiment 2. Once again, the SD was significantly greater for oddballs than standards $t(21) = 2.88$, $p = 0.01$, $d_z = 0.66$ & $t(21) = 2.88$, $p = 0.01$, $d_z = 0.66$. Error bars show ±1 SEM.
